# Melatonin alleviates valproic acid-induced neural tube defects by modulating Src/PI3K/ERK signaling and oxidative stress

**DOI:** 10.1101/2023.07.30.551130

**Authors:** Yuxiang Liang, Ying Wang, Xiao Zhang, Shanshan Jin, Yuqian Guo, Zhaowei Yu, Xinrui Xu, Qizhi Shuai, Zihan Feng, Binghong Chen, Ting Liang, Ruifang Ao, Jianting Li, Juan Zhang, Rui Cao, Hong Zhao, Zhaoyang Chen, Zhizhen Liu, Jun Xie

## Abstract

Neural tube defects (NTDs) represent a developmental disorder of the nervous system that can lead to significant disability in children and impose substantial social burdens. Valproic acid (VPA), a widely prescribed first-line antiepileptic drug for epilepsy and various neurological conditions, has been associated with a fourfold increase in the risk of NTDs when used during pregnancy. Consequently, urgent efforts are required to identify innovative prevention and treatment approaches for VPA-induced NTDs. Studies have demonstrated that the disruption in the delicate balance between cell proliferation and apoptosis is a crucial factor contributing to NTDs induced by VPA. Encouragingly, our current data reveal that melatonin (MT) exerts significant inhibition on apoptosis while promoting the restoration of neuroepithelial cells proliferation impaired by VPA. Moreover, further investigations demonstrate that MT substantially reduces the incidence of neural tube malformations resulting from VPA exposure, primarily achieved by suppressing apoptosis through the modulation of intracellular reactive oxygen species levels. In addition, the Src/PI3K/ERK signaling pathway appears to play a pivotal role in VPA-induced NTDs, with a significant inhibition observed in the affected samples. Notably, MT treatment successfully reinstates the Src/PI3K/ERK signals, thereby offering a potential underlying mechanism for MT’s protective effects against VPA-induced NTDs. In summary, our current study substantiates the considerable protective potential of MT in mitigating VPA-triggered NTDs, thereby offering valuable strategies for the clinical management of VPA-related birth defects.

## 1 Introduction

Neural tube defects (NTDs) is the second most common birth defect in the world and its incidence is second only to congenital heart disease(Mcdonnell *et al*. 2015, Yousif&Mamakan 2022). NTDs are serious birth defects of central nervous system caused by neural tube closure malformations in the early stage of embryonic development, which results in a huge social and economic burden(Zhu *et al*. 2012, Chen *et al*. 2023). NTDs are polygenic genetic disease, which occur under the influence of genetic factors, environmental factors, chemicals, drugs and other factors, although the etiology is not completely clear(Copp *et al*. 2013, Saba *et al*. 2023). The detailed molecular mechanism underlying NTDs remains to be further elucidated(Greene&Copp 2014, Engelhardt *et al*. 2022).

It is reported that, in the United States, about 7.6 million to 12.7 million women suffer from epilepsy each year, and nearly 25,000 of them are pregnant(Meador *et al*. 2008, Harden *et al*. 2009, Li&Meador 2022). Valproic acid (VPA) is a clinical antiepileptic drug and is often used in the treatment of mental disorders such as mania and bipolar disorder, although administering VPA in early pregnancy can lead to a four-fold increase in the incidence of fetal NTD(Lindhout *et al*. 1992, Finnell *et al*. 2021, Zamek - Gliszczynski *et al*. 2022). Given that VPA stands as the most efficacious antiepileptic drug presently accessible, it becomes imperative to diligently observe the preventive measures against NTDs induced by VPA (Duncan 2007). Among the mechanisms through which VPA can impede the normal neural tube development, encompassing the disruption of oxidative stress, DNA hypomethylation, histone deacetylation, and alteration of tetrahydrofolic acid content, it is the perturbation of oxidative stress that assumes a central and pivotal role(Ornoy 2009, Tung&Winn 2011b). VPA induces a surplus generation of reactive oxygen species (ROS) within neuroepithelial cells, triggering apoptosis, and consequently impeding the timely closure of the neural tube. This disruption primarily manifests as malformation with exposed brain tissue. (Tung&Winn 2011b, Hansen *et al*. 2021) The imperative lies in the exploration of efficacious prophylactic interventions aimed at mitigating fetal NTDs resulting from VPA administration.

Melatonin (N-acetyl-5-methoxytryptamine, MT) is an indoleamine synthesized mainly in the pineal gland and is originally found to have physiological and neuroendocrine functions as a diurnal hormone.(Reiter *et al*. 2014, Juhnevica-Radenkova *et al*. 2020) Meanwhile, MT is a clinically reliable antioxidant with strong scavenging effect on free radicals(Naveenkumar *et al*. 2020). The mechanism underlying MT’s resistance to free radicals and oxidative stress involves its direct elimination of free radicals and their byproducts by up-regulating the expression of antioxidant enzymes, restraining the activation of catalase, and preserving mitochondrial homeostasis.(Zhang&Zhang 2014, Fathi *et al*. 2023).

Given the pathogenesis of fetal NTD induced by VPA and MT’s inherent antioxidant capabilities, we postulate that MT exerts a safeguarding influence against the adverse repercussions of VPA administration on NTD occurrence in the clinical context..

In the current investigation, we have employed an animal model of VPA-induced rodent NTD and an in vitro cell model utilizing HT-22 cells to explore the beneficial protective influence of MT against VPA-induced NTD. This comprehensive study delves into both in vivo and in vitro mechanisms underlying MT’s effects. The findings elucidate that MT exerts a corrective influence on the expression of antioxidant genes by modulating the ERK and src signal pathways, thereby ameliorating the heightened fetal oxidative stress resulting from maternal VPA administration. Consequently, this intervention effectively diminishes the occurrence of NTD. These current research outcomes offer novel insights for the clinical prevention and treatment of VPA-induced NTD, holding promising potential in the realm of translational medicine.

## 2 Material and methods

### 2.1 Reagents and antibodies

VPA(P4543), Melatonin (M5250), DCFH-DA (D6883) were ordered from Sigma Chemical Co. (USA). Primary antibodies against src (#2109) p-src (#12432), ERK (#68303), p-ERK (#4370), PI3K (#4249), p-PI3K (#17366) were purchased from cell signaling, PH3 (ab177218), Ki67 (ab16667) and BCL-2 (ab182858) was ordered from Abcam, GAPDH (60004-1-Ig) was purchased from Proteintech Group. FAST DiO (D3898) were purchased from Thermo Fisher Scientific. DMEM/F-12 (11320-033) and fetal bovine serum (10099141) were ordered from Gibco. The ROS Assay Kit was purchased from Beyotime (Shanghai, China).

### 2.2 In vivo mouse experiments

All 6-7-week-old SPF male and female CD-1 mice were raised in the Environmental Barrier Animal Laboratory of Experimental Animal Center of Shanxi Medical University with animal license number: SYXK (Jin) 2019-0007. All animal experiments were examined and approved by the Ethics Committee of Shanxi Medical University. All animal procedures strictly follow the National Institutes of Health guidelines for the Care and use of Experimental Animals.

The mice were maintained on a 12-hour light/dark cycle (lights on from 8:00 am–8:00 pm), with water and food provided ad libitum. Pregnant female mice were obtained by mating with fertile males of the same strain (day 0.5 is the day of the vaginal plug). On day 7.5 of pregnancy (E7.5), VPA was intraperitoneally injected only once at a dose of 400 mg/kg to establish the NTDs embryo model. From E7.5 evening, pregnant mice were given 10 or 20mg/kg of body weight MT (Sigma-Aldrich) by daily i.p. injection until being killed (VPA+MT10 group and VPA+MT20 group). Mice of the control and VPA groups were injected with normal saline in the same manner. The brains of embryos were collected for further analysis.

### 2.3 Analysis of gross morphology

The number of embryo implantation and viable fetuses were carefully examined on E10.5, and the incidence of viable fetuses was calculated as a percentage of the number of implantation sites. Viable fetuses were then screened for NTDs with a stereomicroscope. The incidence of NTDs was calculated as a percentage of viable fetuses.

### 2.4 In vitro cell culture and treatment

HT-22 (Immortalized hippocampal neuron cell), a surrogate of neuroepithelial cell, were cultured in DMEM medium containing 10% FBS. To explore the protective effect of MT on the toxicity effects of VPA, HT-22 cells were co-treated by VPA 10 mM and MT 20 μM for 48h. Cultured cell were harvested for further analysis.

### 2.5 EdU labeling and immunofluorescence staining

Cell proliferation was also analyzed using Cell-Light Edu Kit (RiboBio, Guangzhou, China) following manufacture’s instruction. Immunofluorescence signals were visualized and imaged using a immunofluorescence Microscopy (Nikon, Tokyo, Japan). Image J was adopted to quantified the proliferative cells in different random optical fields.

Immunofluorescence staining of PH3. After being fixed in 4% neutral formaldehyde overnight and embedded in paraffin, fetus brains were cut into 4 μm sections, deparaffinized in xylene, rehydrated through a graded series of ethanol, and washed in water. Then sections were performed antigen retrieval in PH6.0 Sodium citrate buffer and blocked with 10% donkey serum at 37 °C for 1 h. Next, the sections were incubated with primary antibody of PH3 overnight at 4 °C, fluorescent-conjugated secondary antibody for 40 min at room temperature and counterstained with DAPI (1 µg/ml in PBS) for 10 minutes. Images were taken by Nikon ECLIPSE Ti2 fluorescence microscope system. Image J was adopted to quantified the positive signals.

### 2.6 TUNEL assay

TUNEL staining assay was performed as described previously. In Situ Cell Death Detection kit, POD (Roche, USA) was used to assess apoptosis following manufacture’s instruction. HT-22 cell crawling plate or E10.5 embryo paraffin sections were washed three times in PBS and fixed with 4% paraformaldehyde for 30 min at room temperature, then incubated in 3% H_2_O_2_ in methanol in the dark for 20 min. After washing three times with PBS, samples were incubated in a mixture of TdT and dUTP (1:9) from the TUNEL kit at 37°C for 1 h, followed by incubation with converter-POD at 37°C for 30 min. Lastly, samples were counterstained with DAPI for nuclear staining. TUNEL-positive cells were observed under a Fluorescent microscope (Olympus, Tokyo, Japan).

### 2.7 Measurement of ROS

Intracellular ROS level were detected by an ROS Assay Kit (D6883, Sigma) according to the manufacturer’s instructions. Briefly, cells were stained with DCFH-DA (10 µM) and incubated in a cell incubator at 37 °C for 20 minutes after washing three times with serum-free cell culture medium to fully remove the DCFH-DA that did not enter the cells. Cell fluorescence was then detected by fluorescence microscopy (Nikon, Tokyo, Japan) as described by ours(Liang *et al*. 2021).

### 2.8 Western blot

Western blotting was carried out as mentioned previously(Zhang *et al*. 2020). Briefly, total protein of the fetus brain samples was extracted on ice with RIPA lysate and diluted to a uniform concentration. 10 μg protein was loaded on 10% SDS PAGE for separating and transferred to PVDF membranes, which was blocked with 5% skim milk and hybrid with primary antibody. Then, the membrane was incubated in 5% nonfat milk containing HRP-conjugated secondary antibody (1:5000) for 1 h. Signals were detected by an ECL kit (Millipore, USA). Experiments were repeated at least three times. ImageJ was used to detect the gray value of all protein pictures. GAPDH was used as a housekeeping control.

### 2.9 Real-time PCR

Real-time PCR was performed as previously described. Briefly, total RNA was isolated from E10.5 embryos using Trizol reagent kit (Invitrogen); PrimeScript reverse transcriptase reagent kit (R223-01, Vazyme) was used to reverse-transcribe RNA into cDNA. Real-time PCR was performed by using a SYBR Premix Ex Taq kit (Q711-02, Vazyme) on the QuantStudio 5 Real-Time System (ABI, USA). The data were normalized against the levels of *Rpl7* expression. The primer sequences used in this study are listed in Table 1.

### 2.10 Flow cytometry

Flow cytometry was used to analyze the effects of VPA and MT on cell apoptosis with an annexin V-fluorescein isothiocyanate (FITC) apoptosis detection kit (eBioscience, San Diego, CA, USA). After treatment by VPA and MT, HT-22 cells were collected, then centrifuged at 400 × g for 5 min. Cells were grouped into three groups as follows: (1) control group, (2) VPA group, and (3) VPA + MT group; Cells were subsequently incubated with 5 μL of annexin V-FITC and 10 μL of propidium iodide (Sigma-Aldrich, USA) staining solution for 15 min at room temperature in the dark, and cell apoptosis in each group was detected by flow cytometry (BD Biosciences, San Jose, CA, USA).

### 2.11 HE and Immunohistochemistry

Immunohistochemistry (IHC) was performed as described previously (Liang et al., 2018). Briefly, fetus brains were fixed in 4% neutral formaldehyde overnight and embedded in paraffin. 4 μm sections were deparaffinized in xylene, rehydrated through a graded series of ethanol, and washed in water. HE stains kit was used to morphology description. After antigen retrieval, endogenous horseradish peroxidase (HRP) inactivated and blocking, sections were incubated with the indicated primary antibody. Then, DAB Horseradish Peroxidase Color Development Kit was used to detect the positive signals according to the manufacturer’s protocol (Zhongshan Golden Bridge, Beijing, China). The sections were counterstained with hematoxylin.

### 2.12 Statistical analysis

For mouse studies, at least three mice were included in each group. Two datasets were analyzed by Student’s t test. Nonnormally distributed data were analyzed by the Kruskal-Wallis test. The 2^-ΔΔCt^ method was used to analyze PCR results in all experiments. Quantified data were expressed as the mean ± S.E.M. A P value < 0.05 was considered statistically significant. Statistical analysis was performed using GraphPad Prism 6.0 software.

## 3 Results

### 3.1 MT reduces the incidence of VPA-induced abnormal neural tube closure

In order to investigate the preventive effect of MT on the embryonic toxicology of VPA, an NTD mouse embryonic model were established by intrapitoneal injection of 400 mg/kg VPA in CD1 mice, which was comparable to previous studies(Akimova *et al*. 2017, Steele *et al*. 2022). VPA treated embryos displayed an obvious growth retardation and malformations along with a small and hypoplastic brain vesicle by stereomicroscope observation (Figure 1A). Transverse sections were performed on the brain vesicles of fetal mice in each group (Figure 1B), and HE staining showed that fetal forebrain failed to close in VPA treatment group, while the fetal brain vesicles closed completely in the VPA+MT group (Figure 1C). The total incidence of NTDs in this VPA model was up to 53.7%, while the incidence of NTDs was 14% in VPA and MT 20mg/kg co-treatment group (Table and Figure 1D). Those data suggested that neural tube closure was perturbed in VPA treated mouse embryonic tissue, and the incidence of NTD could be reduced by MT significantly (P<0.05)

**Figure 1.**
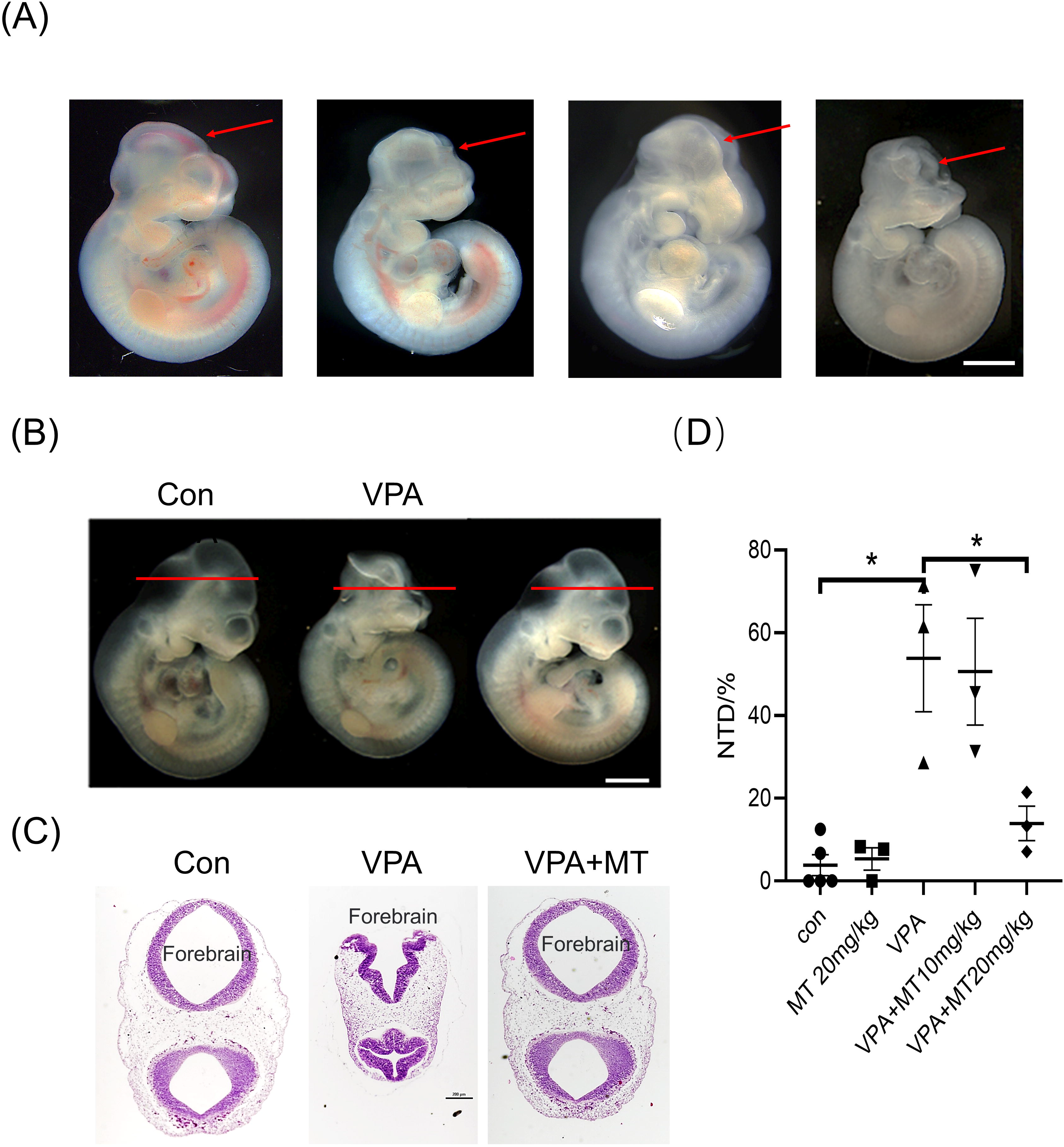
MT reduces the incidence of VPA-induced abnormal neural tube closure. (A) Representative images showing the morphological developmental defects of the brain in VPA treated group. (B) Comparison of fetal brain vesicle morphology and section direction in control group, VPA group and VPA+MT group. (C) HE staining of fetal brain vesicle in control group, VPA group and VPA+MT group. (D) The total incidence of NTDs in control group, VPA group and VPA+MT group. Each group included at least three mice. All data were expressed as means ± S.E.M. *P<0.05, * *P<0.01, ***P<0.001.

### 3.2 MT blocks proliferation inhibition of neuroepithelial cells in the mouse forebrain caused by VPA

During the development of mouse embryonic neural tube, cell proliferation is the basic developmental process to ensure the normal development of neural tube.(Nikolopoulou *et al*. 2017, Lee&Gleeson 2020) The proliferation of neuroepithelial cells in the E10.5 forebrain from all the groups was detected by PH3 and ki67 assay. Compared with the control group, PH3 immunofluorescence signal decreased significantly in the VPA group, while PH3 signal increased significantly in the VPA+MT group compared with the VPA group (Figure 2A). A histogram of the quantitative results of the number of PH3 positive cells was shown in figure2B. Western blot was used to detect the levels of PH3 protein in each group, which was consistent with the results of immunofluorescence (Figure 2C). Quantitative histogram was also used to quantify the results of Western blot (Figure 2D). Moreover, ki67, a classical marker of cell proliferation, was also used to detect the proliferation of anterior neuroepithelial cells in each group by immunohistochemistry. The data showed that compared with the control group, the positive ki67 signal in the VPA group was reduced, while in the VPA and MT co-treatment group, the positive ki67 signal basically recovered to the level of the control group (Figure 2E). The quantitative results of ki67 positive cell number were shown in Figure 2F. This part of data proved that MT could prevent the proliferation inhibition of neuroepithelial cells in the forebrain region of E10.5 mouse fetus caused by VPA.

**Figure 2.**
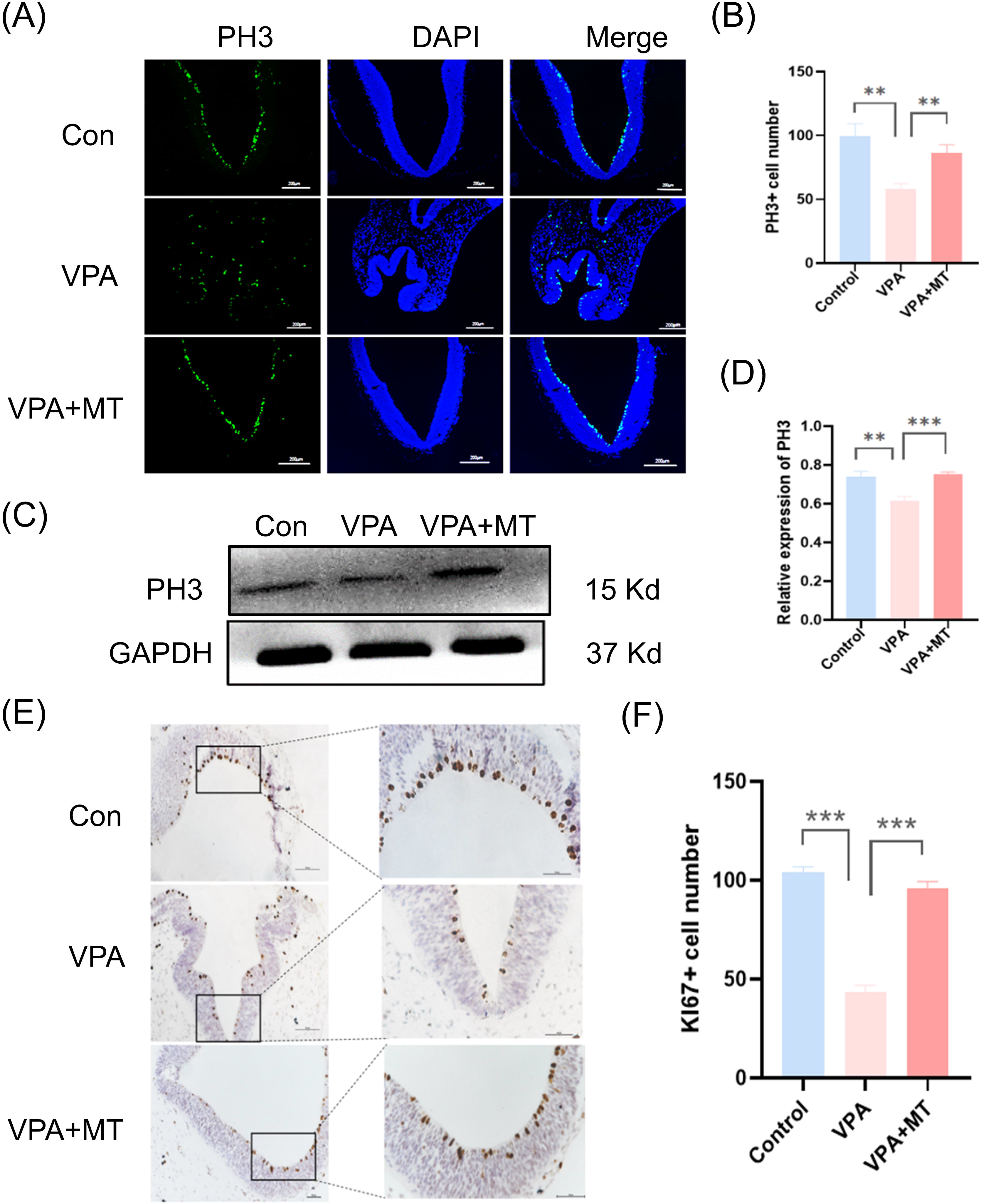
MT blocks proliferation inhibition of neuroepithelial cells in the mouse forebrain caused by VPA. (A) A representative image showing PH3 immunofluorescence of fetal brain vesicle in control group, VPA group and VPA+MT group. (B) The quantification of PH3-positive cells in each group of figure1A. (C) The level of PH3 protein of fetal brain vesicle in control group, VPA group and VPA+MT group by Western blot. (D) Quantitative histogram of gray scale value in each group of figure2C. (E) The expression of Ki67 of fetal brain vesicle in control group, VPA group and VPA+MT group by immunohistochemistry. (F) The quantitative results of ki67 positive cell number in each group of figure2E. Each group included at least three mice. All data were expressed as means ± S.E.M. *P<0.05, * *P<0.01, ***P<0.001.

### 3.3 MT alleviates over-apoptosis of neuroepithelial cells in the mouse forebrain caused by VPA

During the development of mouse embryonic neural tube, excessive apoptosis is also an important mechanism of neural tube defects(Lee&Gleeson 2020). Therefore, the TUNEL assay was used to reveal the level of DNA damage in embryonic neuroepithelial cells. The results showed that the TUNEL signal was significantly increased in the VPA group compared with the control group, while the TUNEL signal was reduced to the control level after MT co-treatment (Figure 3A). The quantitative results of TUNEL were shown in a histogram in Figure 3B. This suggests that the reduction of excessive DNA damage caused by VPA may be one of the reasons why MT can reduce the incidence of neural tube. The ratio of Bcl-2/Bax is one of the important indicators reflecting the level of intracellular apoptosis. Therefore, we detected the protein expression levels of Bcl-2 and Bax in each group, and calculated the ratio of them. The results showed that compared with the control group, the apoptosis level of fetal forebrain neuroepithelial cells in the VPA group was significantly increased, while that in the MT co-treatment group was restored (Figure 3C). The quantitative results of Bcl-2 and Bax were shown in a histogram in Figure 3D. The Bcl-2/Bax ratio is shown in a histogram in Figure 3E. These results suggest that MT may play a protective role by inhibiting excessive apoptosis in VPA treated fetus brain vesicle.

**Figure 3.**
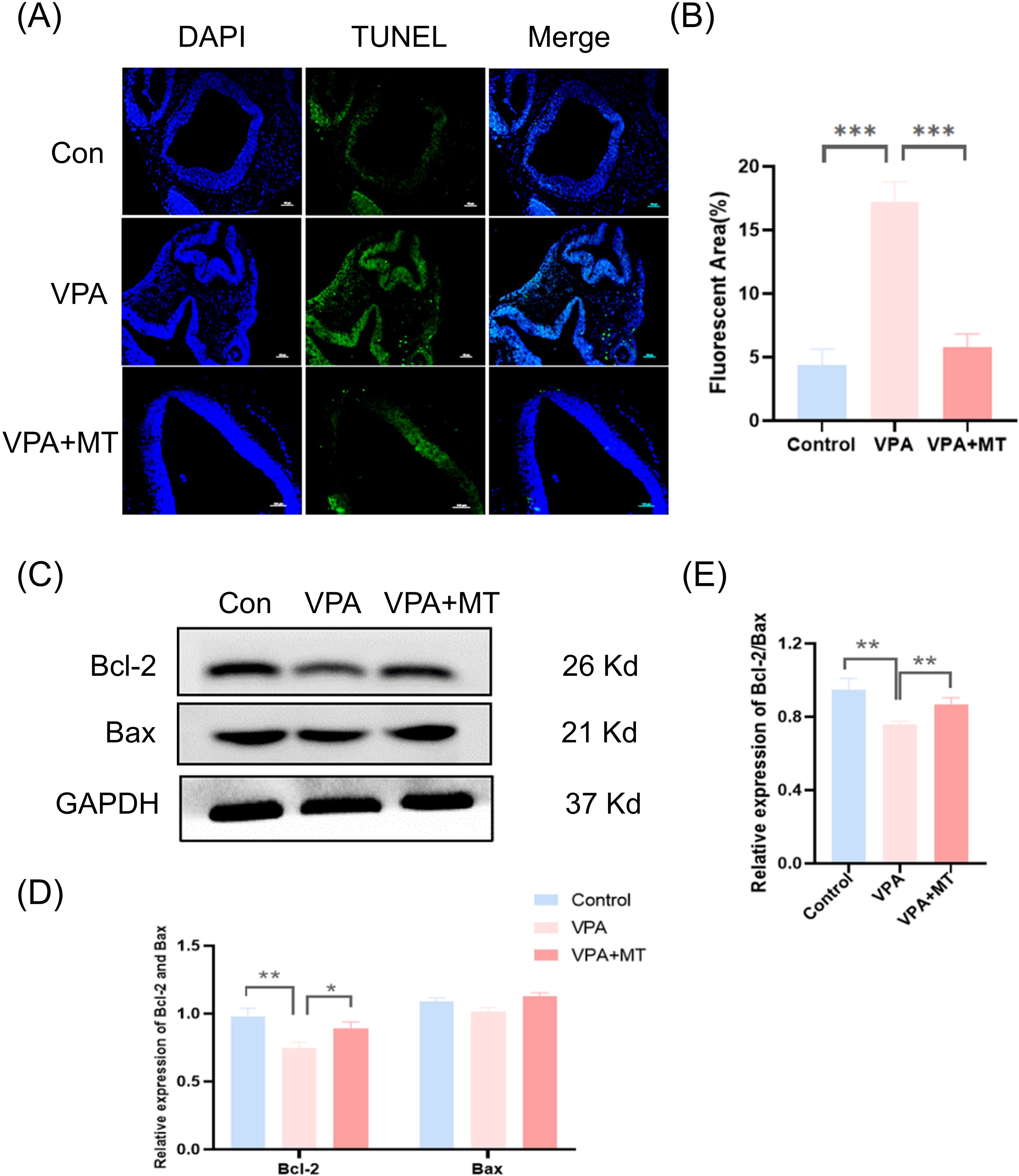
**MT alleviates over-apoptosis of neuroepithelial cells in the mouse forebrain caused by VPA**. (A) TUNEL staining of fetal brain vesicle in control group, VPA group and VPA+MT group. (B) The quantitative results of TUNEL in each group of figure3A. (C) The protein expression levels of Bcl-2 and Bax of fetal brain vesicle in control group, VPA group and VPA+MT group. (D) The quantitative results of Bcl-2 and Bax in each group of figure3C. (E) The Bcl-2/Bax ratio in each group of figure3C. Each group included at least three mice. All data were expressed as means ± S.E.M. *P<0.05, * *P<0.01, ***P<0.001.

### 3.4 MT reverses VPA-induced abnormalities in proliferation and apoptosis of HT22

Further, we investigated the recovery effect of MT on VPA-induced imbalance of cell proliferation and apoptosis in vitro in neuronal cell line HT-22. We conducted EdU staining on the cells in each group, and the results showed that, compared with the control group, VPA could significantly reduce the number of EdU-positive cells, and MT co-treatment could effectively save the proliferation decline caused by VPA (Figure 4A), the histogram of quantitative analysis is shown in Figure 4B. In addition, TUNEL staining of each group of cells was also examined, and it was found that, consistent with in vivo results, VPA significantly induced apoptosis of HT-22 cells, while MT could save the excessive apoptosis caused by VPA (Figure 4C), the histogram of quantitative analysis is shown in Figure 4D. It was necessary to detect the apoptosis level of HT-22 cells in each group by flow cytometry. The results showed that, compared with the control group, VPA induced a large number of apoptosis cells, while MT co-treatment could reduce the number of apoptotic cells (Figures 4E-4H). All in, those in vitro cell results above further corroborate the in vivo findings.

**Figure 4.**
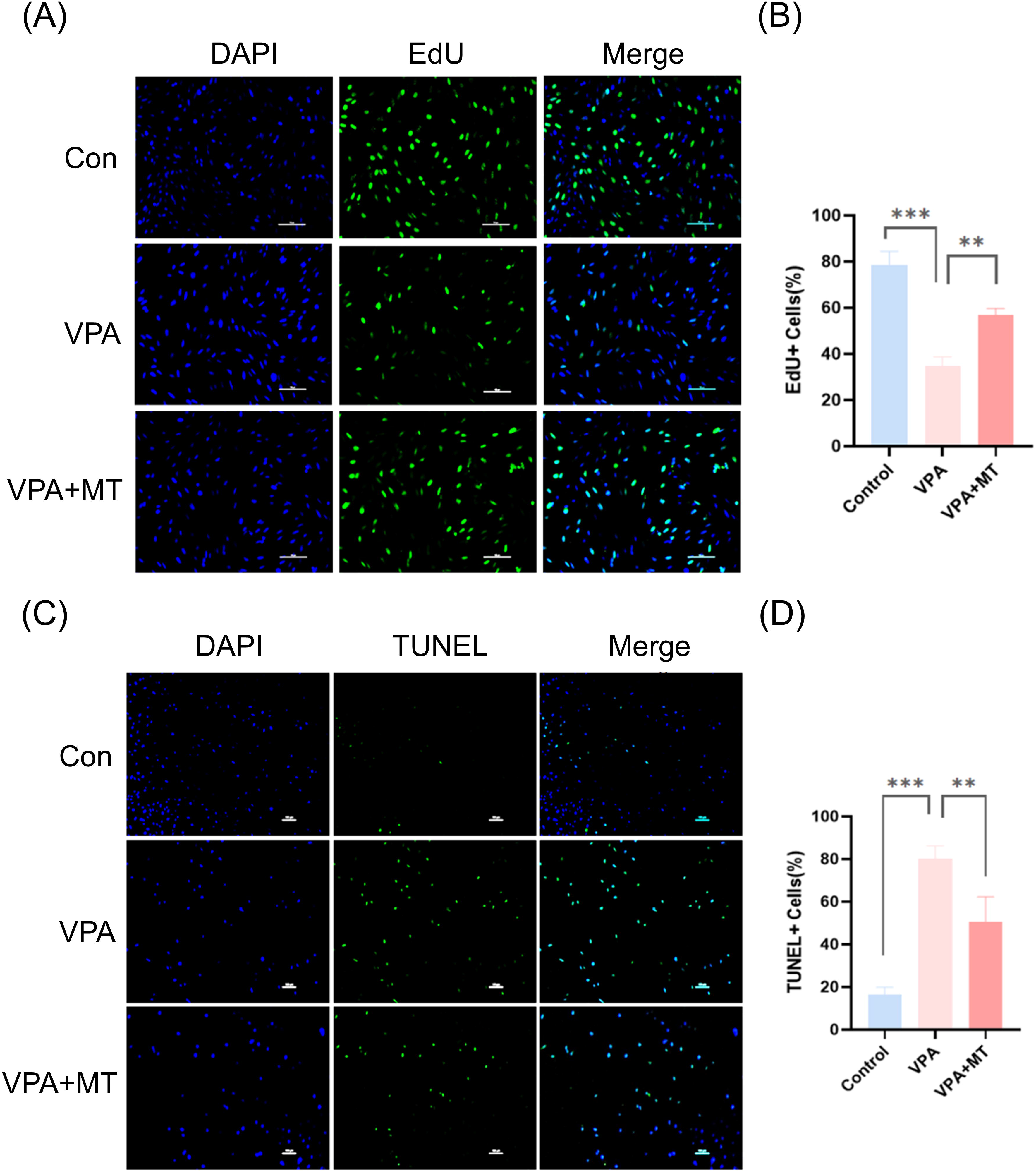
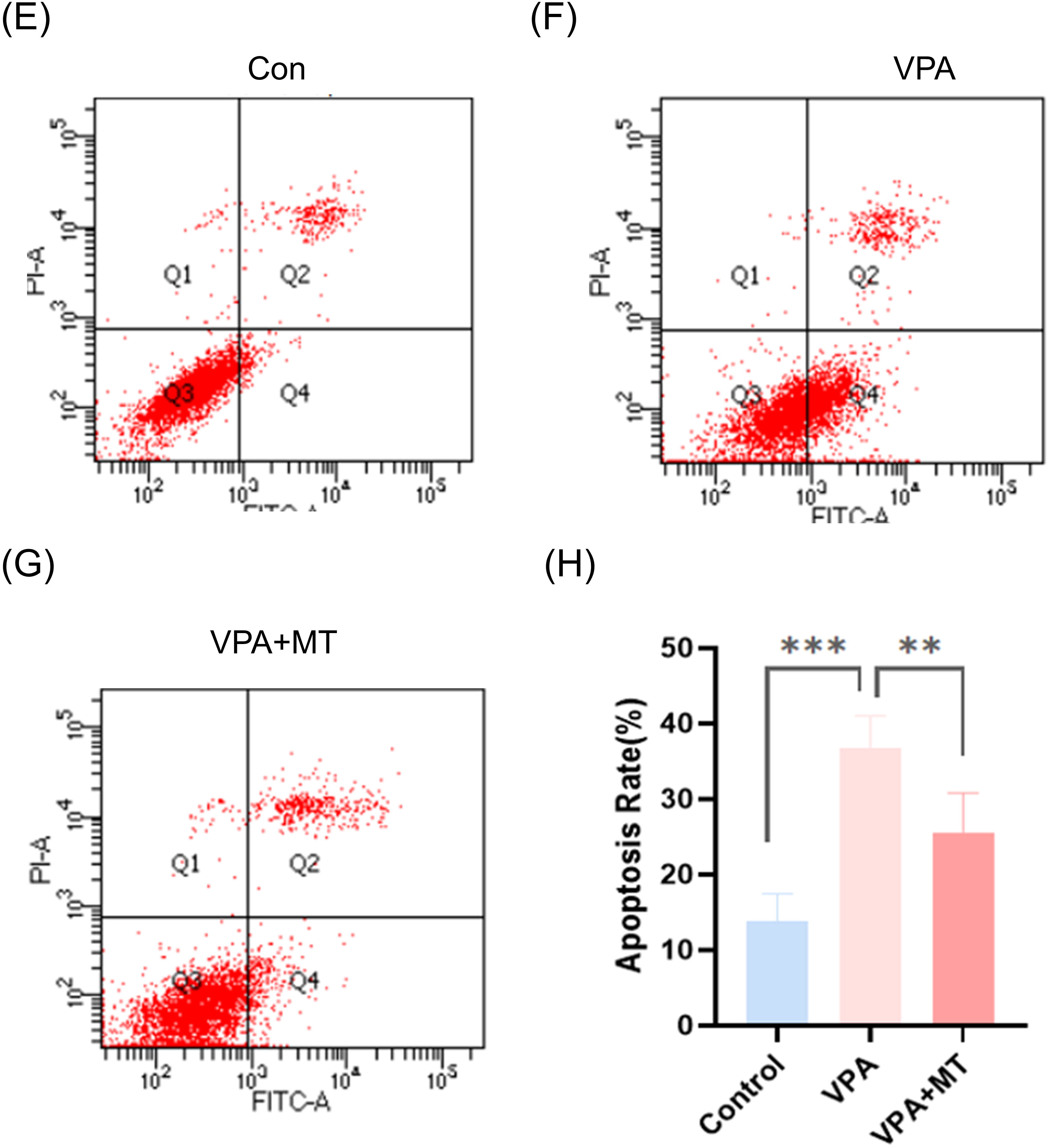
MT reverses VPA-induced abnormalities in proliferation and apoptosis of HT22. (A) EdU staining of HT22 cell in control group, VPA group and VPA+MT group. (B) The quantitative results of EdU in each group of figure4A. (C) TUNEL staining of HT22 cell in control group, VPA group and VPA+MT group. (D) The quantitative results of TUNEL in each group of figure4C. (E-G). Representative image of cell apoptosis in HT22 cell treated by control, VPA and VPA+MT by Flow cytometry. (H) The quantitative plots of apoptosis rates in each group were shown in Figure 4E-G. Each experiment was repeated three times. All data were expressed as means ± S.E.M. *P<0.05, * *P<0.01, ***P<0.001.

### 3.5 MT can alleviate excessive ROS induced by VPA via restoring antioxidant enzymes expression in HT-22 cell

Studies have shown that excessive ROS is an important mechanism of neural tube defects induced by VPA(Tung&Winn 2011b), and excessive ROS production can inhibit cell proliferation and promote cell apoptosis(Diwanji&Bergmann 2018, Luo *et al*. 2019). However, MT is a highly effective antioxidant and free radical scavenger(Gitto *et al*. 2009, Bai *et al*. 2013). We therefore examined the changes in VPA-induced ROS activity following MT treatment on cultured HT-22 for 48 hr. The results showed that compared with the control group, the VPA group had more ROS signal, while MT could significantly inhibit the production of ROS (Figure 5A-5D). In order to further analysis how MT prevent excessive ROS production in fetal brain, gene transcription levels of key antioxidant enzymes in vivo were examined to evaluate the mechanism of MT in reducing ROS. The data showed that compared with the control group, the expression levels of Sod1, Sod2 and gpx1 were down-regulated after VPA treatment, while the expression levels of Sod1, Sod2 and gpx1 were recovered in the VPA+MT group. These results indicate that MT inhibits the production of excess ROS caused by VPA by regulating the expression of antioxidant enzymes in vivo.

**Figure 5.**
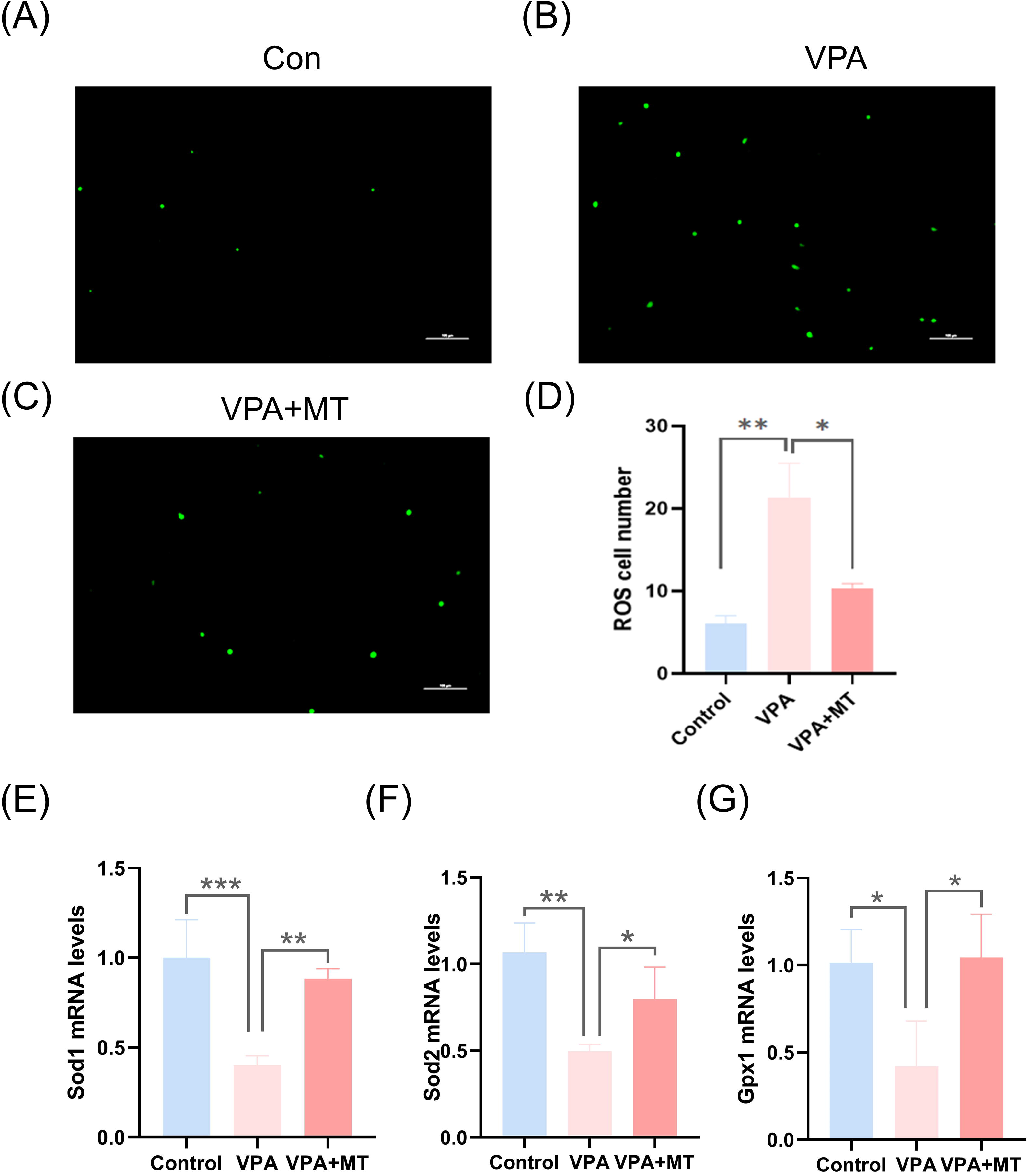
MT can reverse the abnormal proliferation and apoptosis of HT-22 cells caused by excessive ROS induced by VPA. (A-C) Representative image of ROS staining in HT22 cell treated by control, VPA and VPA+MT. (B) The quantitative results of ROS in each group of figure5A-C. (E) The relative mRNA expression of *Sod1.* (F) The relative mRNA expression of *Sod2.* (G). The relative mRNA expression of *Gpx1*. Each experiment was repeated three times. All data were expressed as means ± S.E.M. *P<0.05, * *P<0.01, ***P<0.001.

### 3.6 Src-PI3K-ERK pathway may involve in the preventive therapeutic mechanism of MT against VPA-induced NTD

To elucidate the mechanism by which MT regulates VPA triggered imbalance of cell proliferation and apoptosis in fetus brain, we explored the possible effects of MT on the Src-PI3K-ERK signal transduction pathway. In vivo mouse embryo model, the phosphorylation of src was inhibited by VPA treatment, and MT co-treatment could save the phosphorylation of Src, PI3K and ERK showed similar expression regulation patterns (Figures 6A-D). In vitro cell model, the data showed that MT had rescue effects on p-Src, p-ERK and P-PI3K inhibition induced by VPA to varying degrees (Figures 6E-H), which was mostly consistent with the results of in vivo embryo model. In conclusion, Src-PI3K-ERK signaling pathway may mediate the imbalance of proliferation and apoptosis caused by VPA, while MT may play a role in blocking the occurrence of embryonic neural tube defects by maintaining Src-PI3K-ERK signaling pathway.

**Figure 6.**
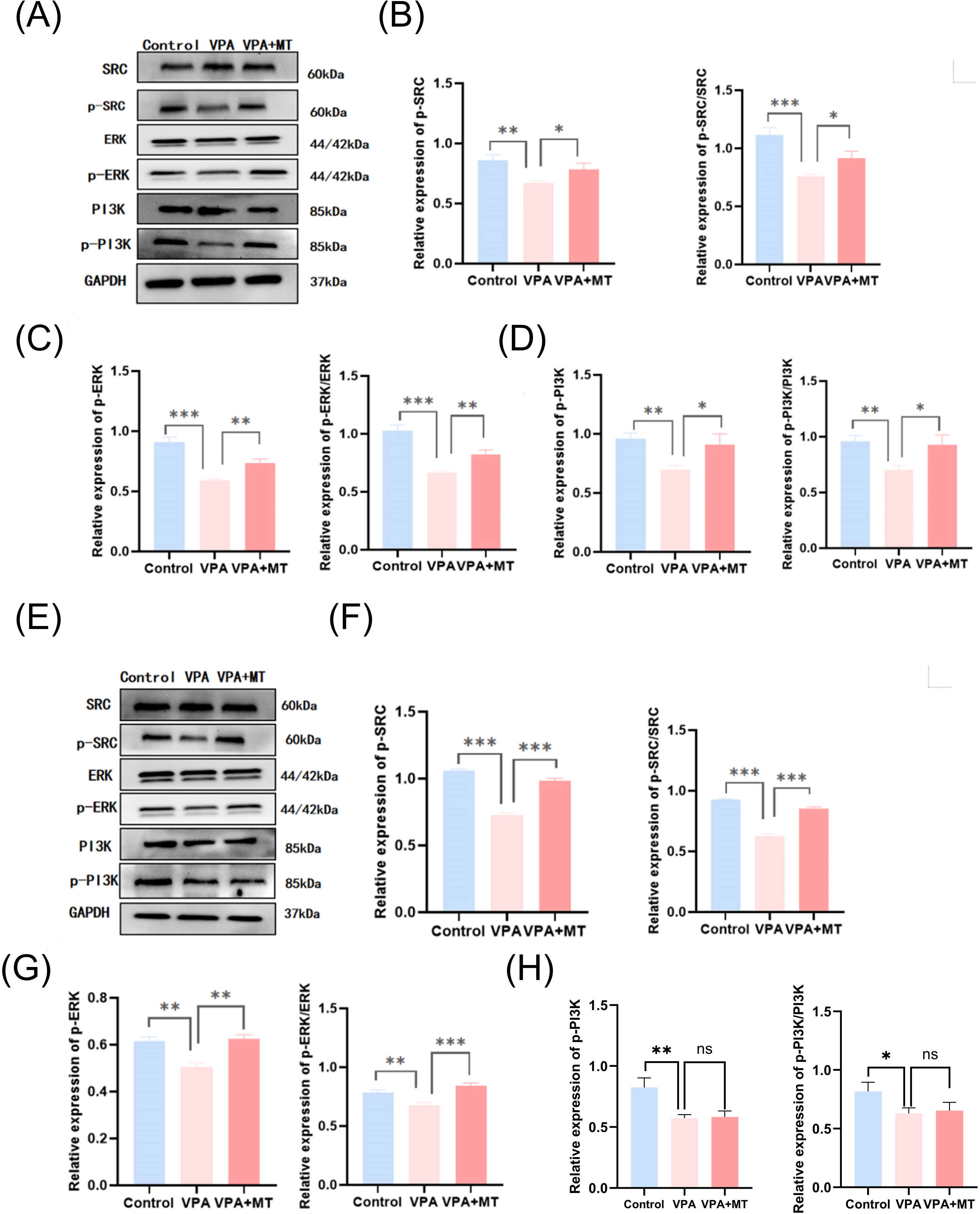
Src-PI3K-ERK pathway may be involved in the preventive therapeutic mechanism of MT against VPA-induced NTD. (A) Western blot analysis of Src/p-Src, PI3K/p-PI3K and ERK/p-ERK levels of fetal brain vesicle in control group, VPA group and VPA+MT group. (B) Quantitative analyses result of Src and p-Src/Src in each group of figure5A. (C) Quantitative analyses result of PI3K and p-PI3K/PI3K in each group of figure5A. (D) Quantitative analyses result of ERK and p-ERK/ERK in each group of figure5A. (E) Western blot analysis of Src/p-Src, PI3K/p-PI3K and ERK/p-ERK levels of HT22 cell in control group, VPA group and VPA+MT group. (F) Quantitative analyses result of Src and p-Src/Src in each group of figure5E. (G) Quantitative analyses result of PI3K and p-PI3K/PI3K in each group of figure5E. (H) Quantitative analyses result of ERK and p-ERK/ERK in each group of figure5E. Each experiment was repeated three times. All data were expressed as means ± S.E.M. *P<0.05, * *P<0.01, ***P<0.001.

## 4 Discussion

The global prevalence of epilepsy affects an estimated 30 million women, with approximately one-third falling within the childbearing age group (Błaszczyk *et al*. 2022). Despite VPA’s recognized teratogenic risks, it remains extensively prescribed for pregnant women with mental disorders. Given that employing VPA in treating epileptic conditions during pregnancy is often an unavoidable necessity(Steele *et al*. 2022), there arises a pressing imperative to innovate and implement preventive approaches that counteract VPA’s teratogenic potential..

The present investigation unveiled that MT effectively mitigates the occurrence of neural tube malformation in a VPA-induced mouse embryonic model. The underlying mechanism appears to involve the inhibition of ROS overproduction and the restoration of PI3K and ERK signaling pathways, culminating in the reduction of neural tube malformation incidence.

While considerable efforts have been directed towards comprehending the neurodevelopmental toxicity mechanism of VPA, a comprehensive understanding of its intricate pathogenesis remains elusive. Nevertheless, a growing body of evidence underscores that a pivotal mechanism contributing to VPA-induced NTDs is the perturbation of redox balance and the subsequent generation of excessive ROS. Emerging research suggests that mitigating redox disruptions could potentially lead to a decline in the prevalence of NTDs. (Tung&Winn 2011b, Hansen *et al*. 2021, Piorczynski *et al*. 2022). Resveratrol and vitamin E could rescue VPA-induced teratogenicity(Hsieh *et al*. 2014). The antecedent study data aligns harmoniously with our current findings, which exemplify that VPA indeed prompts fetal neural tube cells to generate excessive ROS and concurrently inhibits the expression of crucial antioxidant enzymes such as SOD1, SOD2, and GPX1, corroborated by both in vivo animal experiments and in vitro cell experiments.

Melatonin, a hormone synthesized by the mammalian pineal gland, exerts diverse physiological regulatory functions and possesses notable neuroprotective properties. Its capacity to stimulate the proliferation and differentiation of neural stem cells underscores its protective potential against neonatal brain injury induced by hypoxia (Fu *et al*. 2011). Melatonin has demonstrated its protective potential against neural tube defects induced by lipopolysaccharide (Fu *et al*. 2014) and diabetic neural tube lesions in mice(Liu *et al*. 2015). Notably, its favorable lipid-water distribution coefficient facilitates its efficient cellular entry through the cell membrane, enabling the exertion of its potent antioxidant activity(Li *et al*. 2013). In the realm of natural pregnancy, melatonin plays a pivotal role, as plasma melatonin levels in pregnant women rise significantly by 200-300% during the initial 20 weeks of gestation, appearing to be crucial for successful pregnancy outcomes(Tamura *et al*. 2008). Furthermore, the remarkable ability of melatonin to traverse the placenta and cross the blood-brain barrier into the central nervous system positions it as a potential therapeutic agent for preventing VPA-induced NTDs in pregnant women(Liu *et al*. 2015). Despite it is not clear whether MT has a protective effect against VPA-induced NTD until now, our subsequent investigations have shed light on this matter. Encouragingly, our recent findings demonstrate that melatonin effectively diminishes the incidence of VPA-induced NTD by reinstating the expression of essential antioxidant enzymes, namely SOD1, SOD2, and GPX1, and thus counteracting the deleterious overproduction of ROS..

Melatonin therapy is widely acknowledged for its favorable safety profile (Chen *et al*. 2013). Notably, toxicity assessments of melatonin on mouse embryonic development revealed no discernible adverse effects (Mcelhinny *et al*. 1996). Moreover, in a separate investigation, high doses of melatonin administered during pregnancy had no detrimental impact on fetal abnormalities or maternal health (Jahnke *et al*. 1999). Consistent with these findings, our current study demonstrated that the in vivo administration of 20mg/kg melatonin had no discernable effect on the incidence of NTDs.

It has been proposed that the intricate orchestration of neural stem cell processes—proliferation, migration, differentiation, and apoptosis—is a key determinant in neural tube formation(Livingston *et al*. 2023). Studies have substantiated that neural tube disorders can arise from diminished proliferation and heightened apoptosis of neuroepithelial cells within the developing neural tube(Zhao *et al*. 2019, Zhang *et al*. 2021). VPA exposure has been linked to fetal neural tube DNA damage and leading to downstream alterations, including cell cycle arrest and apoptosis(Tung&Winn 2011a). Our findings suggest that melatonin supplementation fosters neuroepithelial cell proliferation and reduces cell apoptosis in VPA-treated fetal brains at E10.5. The results derived from our in vitro HT-22 cell model align with those observed in vivo. An increasing body of evidence supports the notion that oxidative stress can trigger apoptosis (Kannan&Jain 2000, Cao *et al*. 2023), potentially leading to an insufficient number of cells participating in the folding and fusion of neural tube walls.. In this study, melatonin effectively curbed excessive apoptosis of neuroepithelial cells in VPA-treated mice and HT-22 cells in vitro, consequently mitigating the incidence of neural tube malformations. In addition, melatonin treatment significantly elevated the Bcl-2/Bax ratio, decreased TUNEL expression and reduced apoptosis incidence. Notably, melatonin also up-regulate the expression of PH3, KI67 and edU, thus reinstating neuroepithelial cell proliferation. These results collectively highlight melatonin’s antioxidant properties and its role in counteracting apoptosis, aligning consistently with prior research findings(Hong *et al*. 2010).

The Src/PI3K/ERK signaling pathway has been intricately associated with cell proliferation(Yan *et al*. 2019).Inhibition of phosphorylated ERK1/2/MAPK, known for its role in cell proliferation, has been linked to hyperthermia-induced NTDs(Zhang *et al*. 2015). Moreover, considering the pivotal role of PI3K/Akt in embryonic development in embryonic development(Shafique&Winn 2020), its downregulated expression appears to hold significance in the context of VPA-induced neural tube defects. Drawing inspiration from the aforementioned literature, our study sought to elucidate the molecular mechanism underlying MT treatment for VPA-induced neural tube malformations.. Our results unveiled that VPA treatment profoundly suppressed the activation of Src/ERK/PI3K phosphorylation in fetal brain tissue and HT-22 cells. Remarkably, concurrent MT co-treatment significantly restored the levels of Src/ERK/PI3K phosphorylation, suggesting its potential involvement in the amelioration of VPA-induced neural tube malformations by MT. It should be noted that the p-PI3K in HT-22 cell lines treated by VPA and VPA+MT showed no significant difference(figure 6H), although the result of our in vivo experiments in mice (figure 6D) showed that P-PI3K level was significantly restored after MT treatment compared to VPA treatment, so this difference between in vitro and vivo data may not affect the main conclusion of this paper, and the mechanism of the inconsistence need further research in future study.

In conclusion, our current study provides compelling evidence that MT therapy during pregnancy holds promise in averting VPA-induced neural tube disorders by effectively modulating the Src/PI3K/ERK signaling pathway and mitigating oxidative stress. Nonetheless, it is essential to acknowledge that further research is warranted to extend these findings to human applications.

